# Genomic adaptation in group B *Streptococcus* following intrapartum antibiotic prophylaxis and childbirth

**DOI:** 10.1101/2024.04.01.587590

**Authors:** Macy E. Pell, Heather M. Blankenship, Jennifer A. Gaddy, H. Dele Davies, Shannon D. Manning

**Affiliations:** Michigan State University, Department of Microbiology, Genetics, and Immunology (MGI), E. Lansing, MI; Michigan Department of Health and Human Services, Bureau of Laboratories, Division of Infectious Diseases, Lansing, MI; Department of Pathology, Microbiology and Immunology, Vanderbilt University Medical Center, Nashville, TN; Department of Medicine, Vanderbilt University School of Medicine, Nashville, TN; Tennessee Valley Healthcare Systems, Department of Veterans Affairs, Nashville, TN; University of Nebraska Medical Center, Omaha, NE

## Abstract

Through vaginal colonization, GBS causes severe outcomes including neonatal sepsis and meningitis. Although intrapartum antibiotic prophylaxis (IAP) has reduced neonatal disease rates, persistent GBS colonization has been observed in patients following prophylaxis. To determine whether IAP selects for genomic signatures that enhance GBS survival and persistence, a pangenome analysis was performed on 97 isolates from 58 participants before (prenatal) and after (postpartum) IAP/childbirth. Thirty-one of the 34 paired strains from participants with persistent colonization clustered together in the core gene phylogeny, suggesting that the strains recovered at the postpartum sampling were highly similar to those recovered at the prenatal visit. A core-gene mutation analysis identified mutations in 74% (n=23) of the 31 postpartum genomes when compared to the prenatal strains of the same multilocus sequence type recovered from the same individuals. Several strains had acquired mutations in the same colonization-associated genes, though two postpartum strains accounted for most of the mutations. These two outliers were classified as mutators based on high mutation rates and mutations within DNA repair system genes. Increased biofilm production was observed in half of the postpartum strains relative to the prenatal strains, which is supported by the presence of point mutations in genes associated with adherence. Together, these findings suggest that antibiotics may impose a selective pressure on GBS that selects for mutations and phenotypes that promote adaptation and survival *in vivo*. Enhanced survival in the genitourinary tract can lead to persistent colonization, increasing the likelihood of invasive disease in subsequent pregnancies and in newborns following IAP.

## INTRODUCTION

As an opportunistic bacterial pathogen, Group B *Streptococcus* (GBS) colonizes ∼35% of pregnant patients^1^ and colonization is the main risk factor for invasive neonatal infection^2,3^. During pregnancy, rectovaginal colonization can lead to sepsis and other adverse outcomes including preterm birth and stillbirth^4,5^. In neonates, GBS can cause early-onset disease (EOD), which presents as sepsis and/or pneumonia within the first week of life, or late-onset disease (LOD) that presents as bacteremia and/or meningitis between 1 week and 3 months after birth^6^.

In the U.S., screening for vaginal-rectal colonization is recommended between 36 0/7 and 37 6/7 weeks’ gestation^7^. To prevent neonatal infections, GBS-positive patients are given intrapartum antibiotic prophylaxis (IAP) during labor, for which penicillin is the most effective^3^. While IAP has successfully reduced the incidence of EOD, it has not impacted LOD rates nor is it useful for preventing preterm births or stillbirths.^8^ It can also negatively impact the microbiome^9,10^. Importantly, GBS colonization was found to be intermittent during pregnancy with variation in colonization frequencies across individuals and geographic locations^11,12^. One study of EOD cases, for example, showed that 60% of cases were born to individuals whose GBS prenatal screening test was negative^13^, thereby highlighting the need for improved treatment and prevention protocols.

Classifying GBS based on the polysaccharide capsule (cps), which dictates the serotype, and the multilocus sequence type (ST), has identified associations between molecular traits and disease severity^14^. Capsule type III ST-17 strains predominate among neonates with invasive disease^14–16^ and possess unique features that help withstand oxidative stress, survive inside macrophages, associate with host cells, and elicit proinflammatory responses relative to other STs^17–21^. Less definitive epidemiological associations have been observed for other lineages (e.g., STs 19 and 23) despite their high prevalence globally^16^. The effectiveness of IAP was also found to vary across STs in our prior study^22^. Relative to other STs, STs 17 and 19 more commonly colonized patients before and after IAP while persisting for up to six weeks postpartum. ST-12 strains, however, were more frequently eradicated after IAP, which was also independently associated with persistent colonization. Despite these associations, the mechanisms that promote GBS persistence in the vaginal tract and evasion of antibiotic-mediated killing remain elusive. This is a serious concern since persistent colonization enhances the risk of invasive disease, especially for subsequent pregnancies and babies after IAP cessation^10,22^, and may partly explain why GBS LOD rates have not decreased in the U.S.^8,10^.

Altogether, these prior findings led us to hypothesize that IAP imposes a selective pressure on GBS, such that genomic features that promote colonization are selected for, resulting in enhanced fitness and survival in the genitourinary tract. We used whole-genome sequencing (WGS) to compare GBS isolates recovered from pregnant participants sampled both before (prenatal) and 6 weeks after (postpartum) IAP and childbirth^23^. This comprehensive genomics analysis has enhanced understanding of evolution and adaptation among colonizing GBS populations, concurrently providing insight for treatment and prevention improvements.

## RESULTS

### Antibiotic use and GBS colonization dynamics differ across participants

Isolates (n=97) were previously collected^23^ from 58 participants at two time points during pregnancy and after childbirth. The first sampling took place at a prenatal visit between 35-37 weeks gestation, which was the previously recommended timing for GBS screening but differs from recent guidelines of 36-37 weeks gestation^7^. The second sampling took place up to 11 weeks later at a postpartum visit. Among the 58 participants included in this follow-up study, 53 (91.4%) were given IAP during childbirth, with most receiving ampicillin (n=26; 49.1%) or penicillin (n=17; 32.1%) (**Table S1**).

Forty-one of the 58 (70.7%) participants had persistent colonization, meaning they were positive for GBS at both the prenatal and postpartum samplings even though IAP was given to all but three individuals (n=38/41; 92.7%). Among these 41 persistently colonized participants, 35 received a β-lactam antibiotic (ampicillin, n=21; penicillin, n=13; or cefazolin, n=1) and three were given clindamycin. Four of the participants also reported taking antibiotics after childbirth. One was treated with cefazolin and three received either ampicillin (n=1), a combination of trimethoprim and sulfamethoxazole (Septra) with clindamycin and nitrofurantoin (n=1), or metronidazole and cephalexin (n=1) prior to the postpartum sampling. The remaining 17 (29.3%) participants were colonized with GBS only at the prenatal sampling, indicating loss of GBS colonization by the postpartum visit; two of these participants did not receive IAP.

As determined previously^22^, the 97 GBS strains belong to 20 STs comprising clonal complex (CC)-1 (n=25), CC-12 (n=15), CC-17 (n=16), CC-19 (n=17), CC-22 (n=2), CC-23 (n=20), and CC-26 (n=1) plus one singleton (ST-67). Eight distinct cps types were represented, with cps III (n=27) predominating followed by Ia (n=21) and V (n=17); four isolates were non-typeable (NT).

### Pangenome analysis reveals a large accessory genome and clustering by CCs

Ninety-two of the 97 genomes were included in the genomics analysis. Three prenatal and two postpartum genomes from five participants were excluded due to poor assembly quality (**Table S2**), whereas two postpartum strains from two persistently colonized participants (IDs 11 and 14) were not available for WGS. Among the 92 high-quality genomes, 68 represented pairs of strains recovered from 34 participants with persistent colonization (i.e., participants were positive for GBS at both the prenatal and postpartum samplings). Seven unpaired strains from seven persistently colonized participants collected at the prenatal (n=4) or postpartum (n=3) sampling were available for inclusion as well as the 17 prenatal strains from participants who had lost GBS by the postpartum sampling.

The GBS pangenome consists of 5,054 unique genes comprising a small core-genome with 1,368 core and 213 soft-core genes (**Table S3**). It has a large accessory genome containing 945 shell genes and 2,528 cloud genes, which accounts for 68.7% of the total genes. The evolutionary history of all strains was elucidated based on the 1,368 core genes using the Maximum Likelihood method and Tamura-Nei model of nucleotide substitutions in MEGA12,^24^ which produced a phylogenetic tree with the highest log likelihood (**Figure S1**). Five sequence clusters were resolved, grouping together with 100% bootstrap support. Each of the five clusters was associated with a previously defined CC (**Figure 1**). The 17 CC-19 strains and 16 CC-17 strains are grouped into two separate clusters as are the CC-1 (n=23), CC-12 (n=14), and CC- 23 (n=19) strains. Several singletons were detected within each cluster. Construction of a neighbor-net tree in SplitsTree v.19.2,^25^ which was based on 17,825 parsimony-informative (PI) sites within the 1,368 core genes, confirmed the clustering observed in the maximum likelihood phylogeny and uncovered evidence for recombination (pairwise homoplasy index (PHI) p-value = 0.00; **Figure S2**).

**Figure 1:**
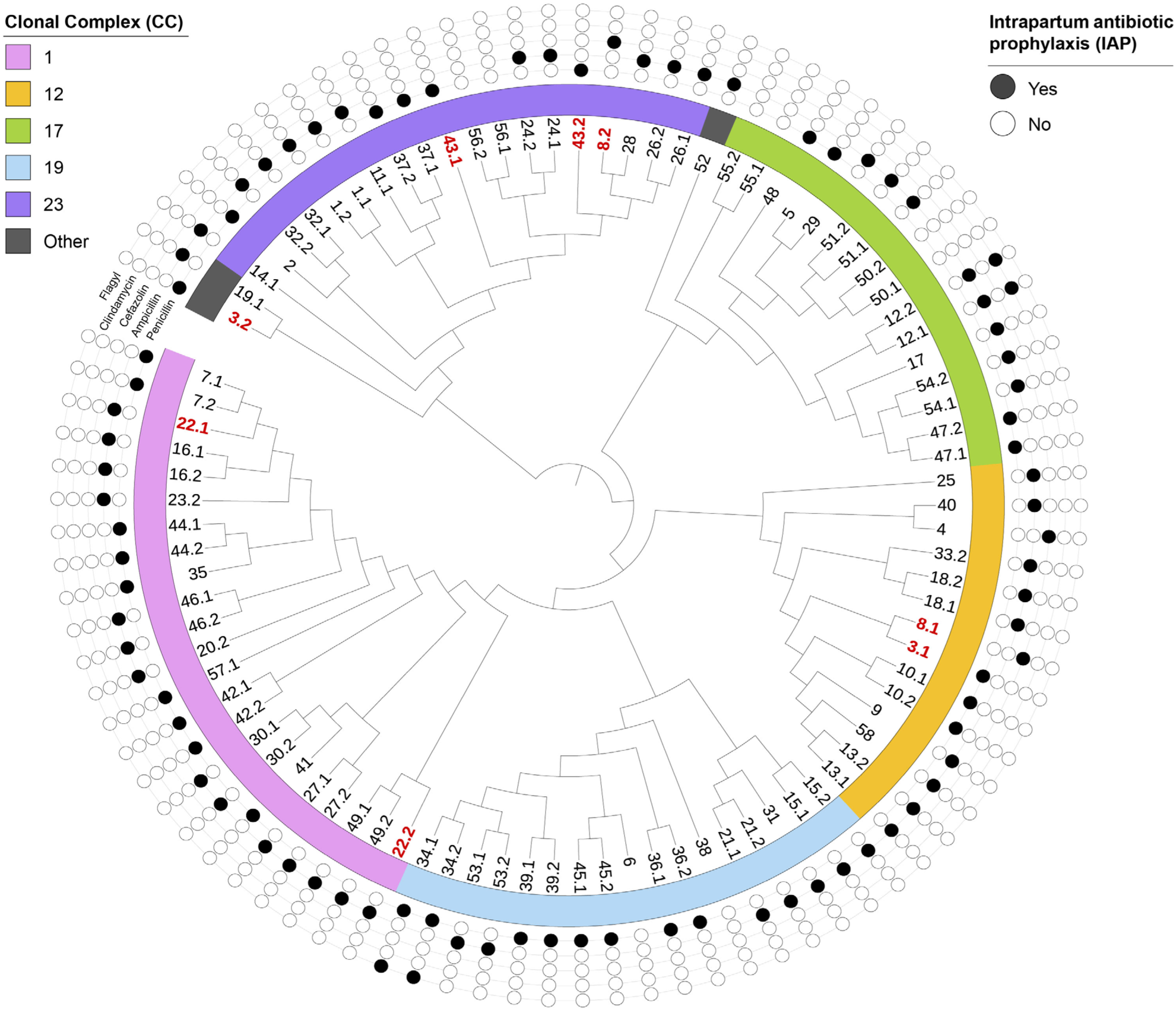
Evolutionary history of 92 GBS strains isolated from participants before and after IAP and childbirth. The bootstrap consensus tree based on 1,368 core genes (1,268,723 positions) was inferred from 88 replicates^26^ with the collapsed branches corresponding to partitions reproduced in less than 50% of replicate trees. The original maximum likelihood tree (Figure S1) represents that with the highest log-likelihood between a Neighbor-Joining (NJ) tree^27^ and Maximum Parsimony (MP) tree. The phylogeny was constructed in MEGA12^24^ and visualized with iTOL^28^ while rooting at the midpoint. Participant strain IDs are noted at the end of each branch; persistent strains contain a “.1” (prenatal) or “.2” (postpartum) indicating the sampling timepoint. The clonal complex (CC) is indicated by colored strips and the circles correspond to the antibiotic(s) used for IAP. Paired strains from the same participant have the same antibiotic usage profiles. IDs in bold red font are the paired prenatal and postpartum strains from four participants (IDs 3, 8, 22, and 43) that do not cluster together in the tree.

Among the 34 pairs of strains from participants with persistent colonization, 30 (88.2%) clustered together in the phylogenetic tree (Figures 1 and S1). The paired strains from the three participants (IDs 50, 55, and 56) who did not receive IAP also grouped together. Of the four pairs that did not group together, two (IDs 3 and 8) had prenatal and postpartum strains with different STs, suggesting acquisition of a new strain type following childbirth and IAP. Although the remaining two participants (IDs 22 and 43) had prenatal and postpartum strains with the same ST, only the prenatal and postpartum strains from participant 43 clustered together. Inclusion of only the paired strains in the consensus tree better illustrates that prenatal strain 43.1 is ancestral to postpartum strain 43.2 (**Figure S3**).

### Acquisition of antibiotic resistance genes is not important for persistent colonization

To determine whether GBS residing in the vaginal tract acquired antibiotic resistance genes (ARGs) to promote colonization, an assembly-based analysis was performed to extract these genes from the 92 genomes. Nine distinct ARGs were identified conferring resistance to tetracyclines (*tetL, tetM, tetO, tetW, lmrP*), macrolides-lincosamides-streptogramins (MLS) (*ermA, lmrP, mreA*), fluoroquinolones (*norB*), and cationic peptides (*mprF*) (**Figure 2**). Both *lmrP* and *mprF* were found in all strains, while *norB* and *mreA* were found in all but one.

**Figure 2:**
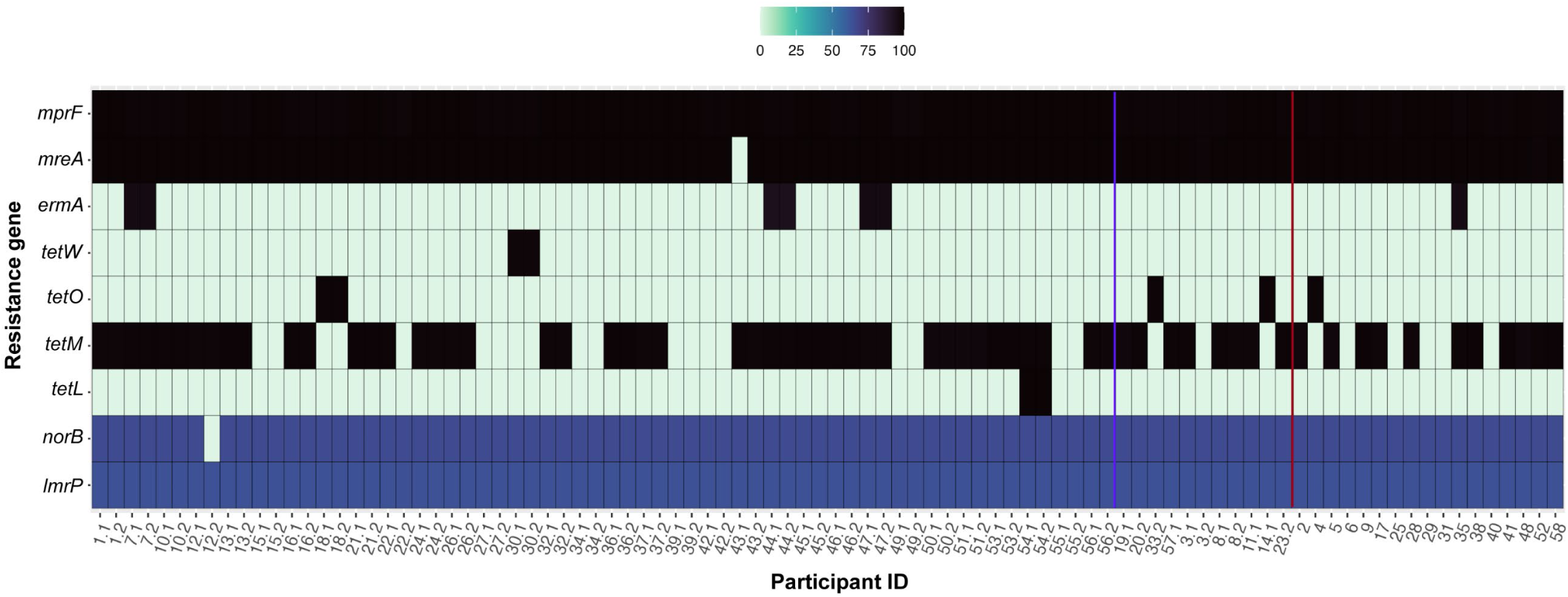
Antibiotic resistance genes (ARGs) detected in the 92 genomes. ARGs were extracted from the 92 high-quality genome assemblies using five different ARG databases. The x-axis shows the patient strains with a “.1” (prenatal) and “.2” (postpartum) after the “Participant ID” indicating the sampling timepoint. The gene profiles for the paired isolates from the same 32 participants with isolates of the same sequence type (ST) are shown next to each other on the left side of the vertical blue line. The two sets of paired strains (IDs 3 and 8) with different STs along with seven unpaired strains are shown between the blue and red vertical lines. The ARG profiles for the 17 “lost” strains, which were recovered only at the prenatal sampling and did not persist, are to the right of the red vertical line. ARGs are indicated on the y-axis that confer resistance to tetracyclines (*tetL, tetM, tetO, tetW, lmrP*), macrolides-lincosamides-streptogramins (*ermA, lmrP, mreA*), fluoroquinolones (*norB*), and cationic peptides (*mprF*). The color gradient shows the average percent identity of a gene across the five databases, with black representing the highest (100%) identity and light green representing the lowest (<10%) identity.

Among the variably present ARGs, *tetM* predominated and was detected in 63 (68.5%) genomes including 47 of the 68 (69.1%) genomes from the 34 participants with persistent colonization. A similar proportion (n=11; 64.7%) was detected in the 17 prenatal strains that were lost following IAP. Although the prenatal and postpartum strains from participant 22 had the same ST, *tetM* was only detected in the prenatal strain (22.1). A discrepancy was also observed between the paired genomes from participant 3, though this finding is less surprising in that the two strains had different STs, strongly suggesting the acquisition of a new strain by the postpartum visit.

### Specific virulence genes or gene profiles are not linked to persistent colonization

The 92 genome assemblies were also interrogated for known virulence genes; 50 distinct genes were detected with 65-100% identity and classified based on function (**Table S4**). Genes that are important for adherence and invasion of host cells (n=18, 36%), immune modulation (n=17, 34%), metabolism (n=1, 2%), toxin production (n=13, 26%), and dissemination (n=1, 2%) were found. The most variable genes were those that are important for capsule (*cps*) production and pili (*pil*) formation (**Figure 3**).

**Figure 3:**
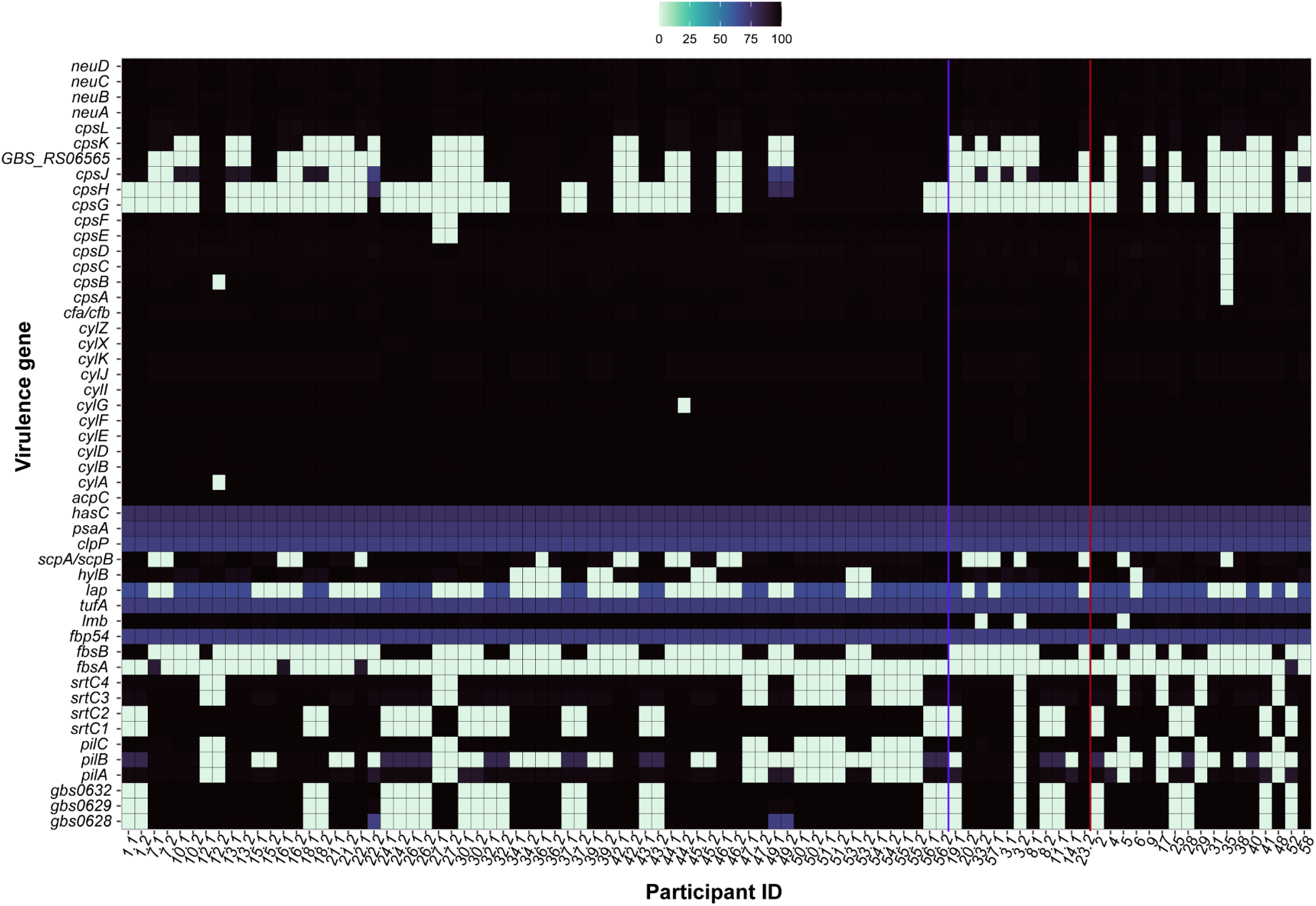
Virulence genes detected in 92 GBS genomes. The x-axis shows the participant ID with the “.1” (prenatal) and “.2” (post-partum) indicating the sampling timepoint. Gene names are displayed on the y-axis. The gene profiles for the paired isolates from 32 participants with isolates of the same sequence type (ST) are next to each other, left of the purple line. The seven unpaired strains plus the two paired strains with different STs from participants 3 and 8 are shown between the purple and red vertical lines. Profiles for the 17 “lost” strains are to the right of the red line. The color gradient displays the average percent identity of a gene, with black representing 100% identity and mint green representing <10% identity.

Among 32 of the 34 participants persistently colonized with strains of the same ST, several discrepancies in gene presence were observed between the prenatal and postpartum strains per pair. In most cases, the differences involved genes with values that fell below the identity and coverage thresholds, indicating that they are present but may be variable. The paired strains from participant 22, however, had multiple differences. Most notably, distinct capsule (cps) types (serotype) were observed in the prenatal (*cpsV*) and postpartum (*cpsVI*) strains, which was demonstrated previously, ^22^ and could indicate capsular switching or acquisition of a new strain by the postpartum sampling. Not surprisingly, participants 3 and 8 with prenatal and postpartum strains belonging to different STs had distinct virulence gene profiles.

To further compare the virulence gene profiles among the paired strains from the participants with persistent colonization, a principal component analysis was performed. No distinct clustering patterns, however, were observed for the prenatal versus postpartum strains (**Figure S4A**) or after stratifying by antibiotic, *cps* type, or colonization phenotype (persistent or lost) (**Figure S4B-E**). In addition, no difference was observed between colonization phenotype and the number of adherence (n=1,031, Chi-square p = 0.10) or immune modulatory (n=1,231, Chi-square p=0.52) genes. These data suggest that neither possession or acquisition of specific combinations of virulence genes were linked to persistent colonization of the vaginal tract after IAP and childbirth.

### The distribution of point mutations varies across persistent strains

To detect nucleotide-level variation, we conducted a reads-based analysis of the 1,368 core genes in the 64 paired prenatal and postpartum strains that shared STs. In this analysis, each prenatal genome was used as the reference genome for its paired postpartum genome and the number and type of mutations in the postpartum genomes were determined. Alarmingly, 7,025 mutations were identified in 24 (75.0%) of the 32 postpartum genomes, ranging from 1 to 6,564 mutations per genome with an average of 292 (**Table S5**). No mutations were detected in eight postpartum genomes, which have identical core gene sequences as their paired prenatal genomes despite IAP exposure. Of the three participants who did not receive IAP, the postpartum isolates had only 2, 3, or 4 total mutations when compared to their respective prenatal isolates.

Among the 24 postpartum genomes with at least one mutation, a skewed distribution of mutations was observed with three outliers (**Figure 4A**). These outlier genomes contained 204, 262, and 6,454 total mutations as opposed to an average of five mutations per genome and range of 1-16 mutations for the remaining 21 postpartum genomes (**Figure 4B**). Most (n=6,065; 86.3%) of the mutations were in coding regions, though an additional 958 (13.6%) were in uncharacterized regions and two were in rRNAs (**Table S6**). The type of mutations varied across genomes (**Figure S5A**) with the majority (n=5,982; 85.2%) representing single nucleotide polymorphisms (SNPs) including synonymous (n=3,469; 58.0%), missense (n=1,771; 29.6%), or nonsense (n=26; 0.4%) mutations (**Figure S5B**).

**Figure 4.**
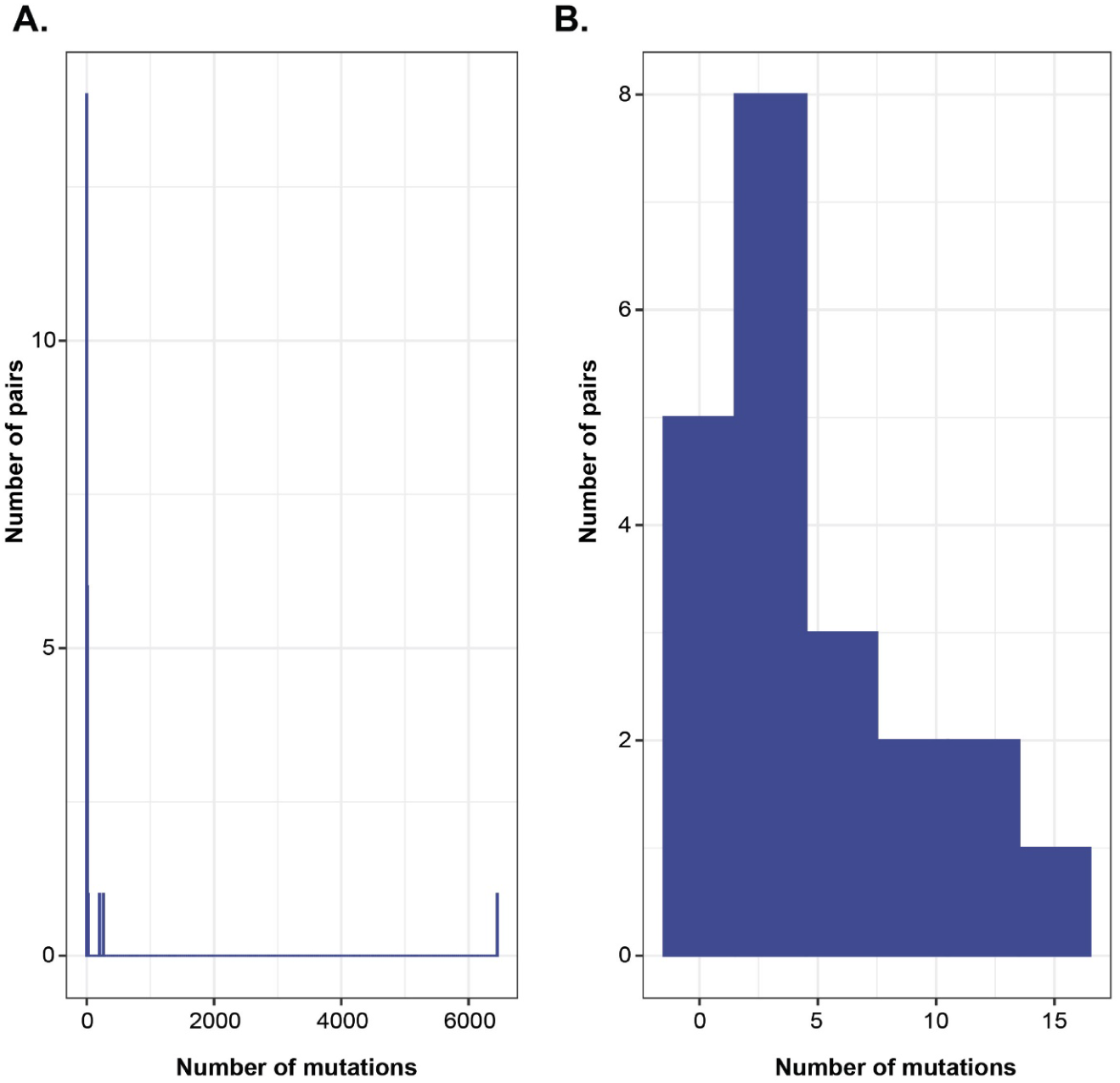
Number of core gene mutations in the postpartum GBS genomes with at least one mutation. Both plots show the number of mutations (x-axis) by the number of genomes (y-axis) for **A)** all 24 postpartum genomes, using a bin width, or interval size, of 10; and **B**) the subset of 21 postpartum genomes after removing three outliers with the greatest number of mutations with a bin width of 3. The eight genomes without any mutations were omitted from these plots.

To define which genes possessed SNPs within coding sequences, we mapped the 5,291 characterized SNPs to specific genes. Most of these SNPs (n=5,233; 98.9%) were detected in the three outlier genomes, mapping to 1,167 genes and yielding an average of 2 SNPs/gene (**Figure S6A**). After excluding all three outliers from the analysis, 58 SNPs were found within 56 genes in the remaining genomes (**Figure S6B**). Within the outlier genomes, most genes (n=683) acquired less than four SNPs relative to the respective prenatal genomes with 390, 142, 86, and 65 genes having 1, 2, 3, and 4 SNPs, respectively. Four or more SNPs, however, were detected within 366 genes. Among these, 91 genes had 662 SNPs, which were mostly classified (98.3%, n=651) as missense (nonsynonymous) mutations. A subset of these missense mutations was found in the same genes in more than one of the three outlier genomes and encode for proteins involved in type VII and Sec-system secretion as well as pilus assembly (**Figure S6C**). Although most (77.6%, n=284) of these 366 genes have evidence of negative selection with dN/dS values < 1.0, 61 (16.7%) have evidence of positive selection with dN/dS values > 1.0 (**Table S7**). Despite the range in type and distribution of mutations that could be classified, the high frequency of mutations within postpartum strain 22.2 skewed the distribution of each type of mutation identified.

### Increased mutation rates and mismatch repair mutations uncover “mutator” strains

Given the wide range and frequency of mutations observed, we calculated the mutation rates based on the total assembly length per genome for comparison. The mutation rate, which represents the number of mutations per number of base pairs (bp), was determined for each postpartum genome assembly relative to its respective paired prenatal genome assembly.

Notably, the average mutation rates for the three outlier genomes were 9.37×10^-5^ (ID=36.2), 1.32×10^-4^ (ID=43.2) and 0.003 (ID=22.2) mutations/bp, which is 53-, 75-, and 1,747-fold higher, respectively, than the average mutation rate (1.77×10^-6^ mutations/bp) for the rest of the 29 genomes including those that lacked any detectable mutations (**Table S5**).

Among the three persistently colonized participants (IDs 50, 55, and 56) who did not receive IAP, the average mutation rate was 1.50×10^-6^ mutations/bp, which is 77.3 times lower than the average rate (1.16×10^-4^) for the 29 participants who were given IAP, and 6.5 times lower after omitting the outlier postpartum genome (22.2) with 6,454 mutations. Although the difference in means was not significant (Student’s t-test p=0.13), the mutation rate for strains without IAP exposure includes the postpartum strain from participant 56 (n=4 mutations), who was one of the four individuals reporting antibiotic use (cefazolin) after childbirth. Nonetheless, these results demonstrate variable mutation rates across strains and suggest that the higher-than-average mutation rates observed in the three outlier postpartum genomes could be indicative of “mutators” that arose following IAP and childbirth.

Indeed, the maximum likelihood core gene phylogeny provides support for mutators in two of the three postpartum strains (Figure 1). Except for strain 22.2, which had the greatest number of mutations, the other two outlier strains, 36.2 and 43.2, grouped together with their respective prenatal genomes. The pair of strains from participant 36 share a node in the phylogeny, while the prenatal strain 43.1 is on an adjacent ancestral branch to its paired postpartum strain 43.2.

These data suggest that the genetic backbones of these two outlier postpartum genomes are highly similar to their respective prenatal genomes despite the high number (>200) of core gene mutations detected. By contrast, the placement of the 22.2 postpartum genome in the phylogeny relative to the prenatal 22.1 genome along with different resistance and virulence gene profiles (Figures 2 and 3), and an unusually large number of mutations, suggests that strain 22.2 is not a mutator. Rather, these data indicate that participant 22 likely acquired a new strain of the same CC with a distinct *cps* type by the postpartum visit.

Since mutators arise at a low frequency in the population due to mutations within genes encoding DNA replication and proofreading machinery, mismatch repair (MMR) pathways, and the 8-oxo-dG (GO) system^29,30^, we compared the genes involved in these pathways among the prenatal and potential mutator strains from participants 36 and 43. The six MMR pathway genes (7,467 bp region) were extracted from each genome and aligned using MUSCLE^31^. These genes encode DNA mismatch repair proteins, MutS (*mutS*) and MutL (GBS_230), a cold shock protein, an MFS transporter/LmrP homolog (GBS_436), the Holliday junction branch migration protein (RuvA), and DNA-3-methyladenine glycosylase I (GBS_235). The postpartum 36.2 genome has a 33 bp insertion in *mutS* relative to its paired 36.1 prenatal genome as well as a deletion in the DNA polymerase IV gene and a missense SNP in *mutY* (A/G-specific adenine glycosylase) of the GO system. Comparatively, genome 43.2 has a missense SNP in the DNA topoisomerase gene and a 1 bp deletion downstream of the MMR loci relative to its paired 43.1 strain. The MMR loci were identical in the remainder of the paired prenatal and postpartum genomes. The increased mutation rates along with mutations in these key regions meet the criteria^29,32^ for classifying these two strains as “mutators”.

### Specific lineages are more likely to acquire mutations *in vivo*

To better understand which point mutations could alter protein functionality, we examined the distribution of the 350 characterized SNPs in the 31 postpartum strains after omitting outlier strain 22.2 from the analysis. While specific mutations were not associated with individual lineages, CCs 19 (n=121) and 23 (n=199) had the greatest number of SNPs, though these numbers are skewed by inclusion of the two mutator strains. After excluding the mutators from the analysis, the CC-23 strains still had the most SNPs (n=21) followed by CCs 12 (n=15) and 1 (n=12), ranging from 0-10, 0-9, and 0-7 SNPs per strain, respectively (**Table S8**). By comparison, the postpartum non-mutator strains of CCs 17 and 19 acquired fewer SNPs overall with 0-1 and 0-3 SNPs per strain, respectively. Among the three SNPs identified in the six CC- 17 postpartum strains, only one was nonsynonymous, indicating a pattern of negative or purifying selection for strains in this lineage. The non-mutator strains of CCs 23, 12, and 1, however, had twice as many nonsynonymous (n=32) SNPs as synonymous (n=16) SNPs, suggesting a pattern of positive/diversifying selection and a trend of adaptive evolution for these lineages.

Based on prior medical record reviews^23^, the two mutator strains had been exposed to penicillin, which was used for IAP in participants 36 and 43 during childbirth (**Table S8**). Among the 16 participants given ampicillin and the 10 receiving penicillin, 28 and 310 SNPs were detected in the postpartum strains, respectively. After excluding the two mutators, the number of SNPs detected among the 8 participants who received penicillin decreased from 310 to 18 SNPs. While comparisons could not be made for the other antibiotics due to low numbers, the three genomes from patients who did not receive IAP only had 1, 3, and 4 total mutations with an average of 1 SNP/strain. The strain with 4 mutations, however, was exposed to cefazolin after childbirth and prior to the postpartum sampling. Collectively, these data demonstrate that certain strain types may be more susceptible to evolutionary pressures, which likely varies depending on the antibiotic used for IAP.

### Persistent strains have enhanced biofilm production relative to prenatal strains

Because biofilm formation is a common colonization strategy and mutations were found in genes important for adherence, we examined this phenotype in the paired strains. An increase in biofilm production was observed in slightly over half (54.8%, n=17) of the 31 postpartum strains relative to each paired prenatal strain (**Figure 5**); four of these increases were significant. Among the two mutators, one (36.2) had increased biofilm production and one (43.2) had decreased production relative to the respective prenatal strains (**Figure S7**), though the differences between both pairs were not significant. It is noteworthy that two postpartum isolates from participants 26 and 27 had a significant decrease in biofilm formation relative to the prenatal isolates. These isolates had acquired 16 and 2 mutations, respectively, with some occurring in genes encoding binding and surface proteins such as penicillin-binding protein Pbp2B, capsule protein CpsC, and an LPXTG anchor domain surface protein.

**Figure 5.**
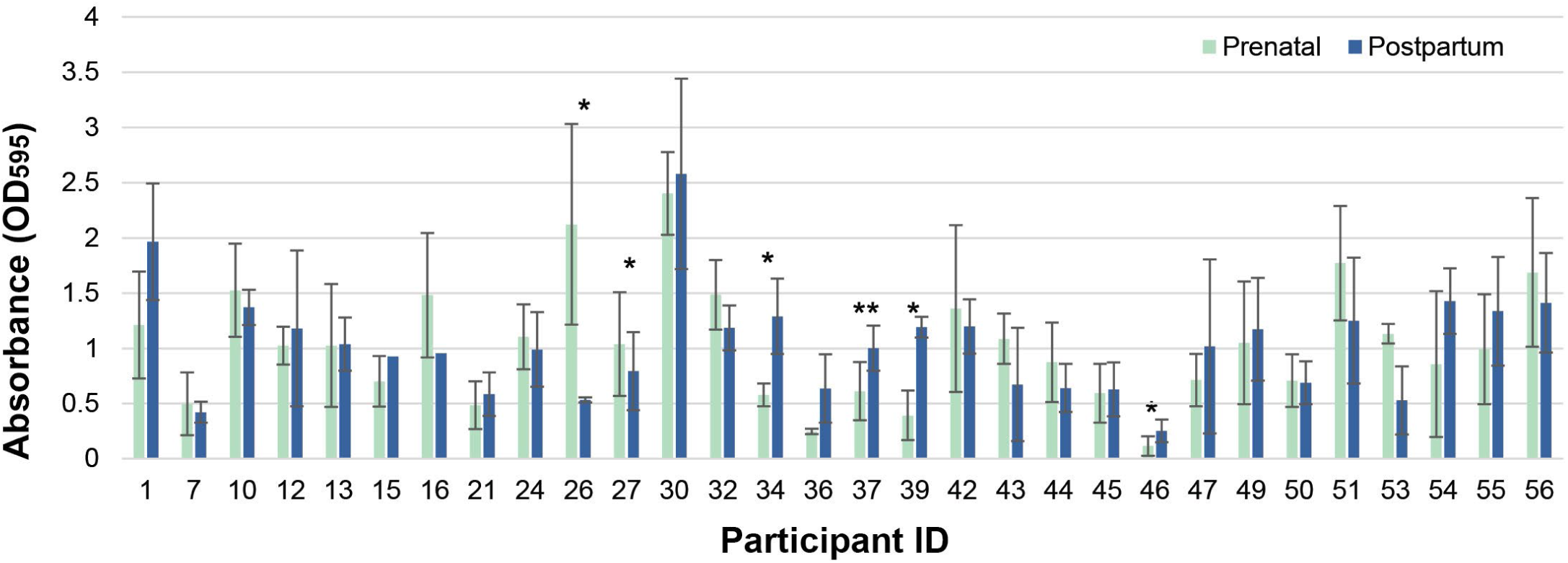
Biofilm formation between prenatal and postpartum strains from 30 participants with persistent colonization. Biofilm formation was assessed across 30 paired strains by measuring the absorbance (OD_595_) values (y-axis) and averaging the biological replicates (n=3). Prenatal (mint green) and postpartum (dark blue) isolates are shown by participant ID (x-axis). One-tailed, paired t-tests were calculated to identify significant differences in biofilm formation between the paired strains (*=p <0.05, **=p<0.01). Each measurement was normalized to a negative media control. Error bars display standard deviation.

## DISCUSSION

Through this analysis, we have identified a range of genomic alterations in GBS strains recovered after IAP and childbirth as compared to strains isolated during prenatal visits from the same individuals. Although the pangenome analysis of 1,368 core genes uncovered five distinct sequence clusters, most of the paired prenatal and postpartum strains with identical STs clustered together in the phylogeny. This finding along with a comprehensive evaluation of point mutations indicates that in most cases, the strains that persisted in each participant are highly similar to those recovered prior to IAP and childbirth.

Although various antibiotic resistance mechanisms are known to enhance bacterial survival in the presence of antibiotics, our assembly-based genomic analysis failed to detect newly acquired resistance genes in the postpartum strains relative to each respective prenatal strain. Hence, resistance to the antibiotics used for IAP is not a mechanism of persistent colonization in this study population. Among the paired postpartum genomes from the 26 (83.9%) participants who received β-lactam antibiotics for IAP, none had β-lactam resistance genes, which is not surprising since GBS remains universally susceptible to β-lactam antibiotics^10^. Resistance to MLS antibiotics is more common in GBS^10^, yet the three (9.7%%) participants given clindamycin for IAP had postpartum strains that were neither phenotypically resistant^23^ or possessed macrolide resistance genes. In a similar analysis, acquisition of specific virulence genes was also not associated with persistence since distinct variably present genes were not detected in the postpartum versus prenatal strains. While these findings indicate that horizontally acquired resistance and virulence genes are not a mechanism of GBS adaptation in this patient population, it is important to note that there are limitations associated with assembly-based analyses. Genes labeled as absent, for instance, may have been present below the detection threshold due to poor coverage or allelic variation, or may exist in a region that is more difficult to sequence. Since the only discrepancies observed were in three pairs of strains that did not cluster together in the core gene phylogeny (Figure 1), we expect that the determination of presence versus absence of each gene was accurate for the remaining pairs.

The accumulation of point mutations appears to be more important for GBS adaptation than horizontal gene transfer in this study population, as was indicated in our reads-based analysis of the 1,368 core genes. Genomic mutations were identified in 74% (n=23) of the 31 postpartum genomes that shared an ST and clustered together with their respective prenatal genomes in the core gene phylogeny (Figure 1). While this core gene mutation analysis has limitations including the possibility of sequencing errors, each mutation was manually confirmed to rule out sequencing artifacts and was determined to be in an area of >10x read coverage. In all, 0 to 262 mutations were detected in the 31 postpartum genomes, though two genomes are outliers that harbor 81.6% (n=466) of the 571 mutations detected (Table S5). Exclusion of these two outliers uncovered 105 mutations in 29 genomes with a range of 0 to 16 mutations per genome. This number is slightly larger than the total number of mutations (n=29) detected among paired strains from 19 pregnant patients and their sick neonates in a smaller WGS study^33^. In the paired neonatal strains, mutations were detected within *covR*, which encodes a two-component response regulator linked to virulence^34^, and other virulence-associated genes. Although *covR* mutations were not observed in our postpartum genomes, several strains had mutations in genes encoding transcriptional regulators and ABC-transporters as well as colonization factors like type VII secretion proteins (*essC*)^35^, serine-rich repeat (srr) glycoprotein adhesins^36^, and sortases (*srtC)* used for pilus formation^37^. It is therefore possible that these mutations enhance fitness *in vivo* while promoting adaptation to the genitourinary tract. Although epistasis, or the impact of bacterial genotype on the fitness effects of a mutation, has been described^38^ and CCs commonly linked to maternal colonization^16^ had the most mutations (Table S8), future experiments are needed to confirm the impact of each mutation on GBS fitness and survival in different strain backgrounds.

Even though we did not test the functional impact of all mutations identified, we did observe functional changes in biofilm production that could be linked to the mutation data. Despite the wide range in biofilm production, which is consistent with studies showing differences across lineages, pilus types^39^, and culture conditions^40^, half of the postpartum strains had an increase in biofilm formation relative to the prenatal strains. In some strains with enhanced biofilm production, point mutations were identified in genes linked to adherence or colonization. Strain 36.2, for instance, had mutations in *srtC* (class C sortase), Sec-system proteins (GtfA, Asp2), and a serine-rich glycoprotein adhesin (*srr1*), which was associated with persistent vaginal colonization in mice^41^. Mutations were also discovered in some isolates with decreased biofilm production. The postpartum isolate with the largest significant decrease in biofilm production has a missense SNP in the capsular polysaccharide biosynthesis protein gene, *cpsC*. Given the tradeoff between capsule production and adherence, which has been described in *S. pneumoniae*,^42^ it is reasonable to speculate that this mutation may promote capsule production while simultaneously blocking adhesins that may be required for aggregation. Additional mutagenesis and functional studies as well as use of more sophisticated biofilm quantification methods, however, are needed to confirm the impact of this and other mutations

The high frequency of mutations observed in the two outlier postpartum genomes, which have 204 and 262 mutations relative to their respective prenatal genomes, led to their classification as mutators. Support for this classification comes from the placement of each pair in the core gene phylogeny (Figure 1) as well as the increased mutation rates in both postpartum genomes and the identification of mutations in genes encoding DNA replication and proofreading machinery^29,32^. Mutators represent a subpopulation of bacterial cells with a higher-than-average mutation rate^29^ that typically arise in populations undergoing adaptation to stressful conditions^43,44^. They serve as a reservoir of mutations that can enhance fitness along with the genetic diversity of bacterial populations^46^. A study of *S. pneumoniae*, for instance, described increased mutation rates after exposure to subinhibitory concentrations of penicillin^45^, which facilitated adaptation. While mutators have been described in *S. pneumoniae*^47^ and other streptococcal species (e.g., *Streptococcus pyogenes* ^48^, *S. mutans*^30^, and *S. iniae*^49^), this study is the first to describe mutators in GBS.

Importantly, mutators have been linked to persistent bacterial infections, such as those caused by *Pseudomonas aeruginosa*^50^ and *Staphylococcus aureus*^51^ in the lungs of cystic fibrosis (CF) patients. They arise at low frequencies due to DNA polymerase errors, dysfunctional proofreading mechanisms, and/or failure of mismatch/error correcting machinery, including the MMR pathway^29,32^. A prior study of pathogenic *Escherichia coli* and *Salmonella enterica*, for example, described the incidence of mutators with defective *mutS* alleles to be 1%^52^; MutS is a critical protein that recognizes mismatches^53^. Impairment of genes in the MMR pathway can contribute to increased mutation rates, recombination,^52^ and horizontal gene transfer, which can subsequently enhance the spread of antibiotic resistance determinants^32^. In *S. aureus*, hypermutable strains recovered from the CF lung had high frequencies of mutations in macrolide target sites^51^, highlighting their role in the emergence of macrolide resistance.

Besides promoting adaptation in the presence of antibiotics, many of the *S. aureus* hypermutators also had mutations in *mutS*. Herein, both GBS mutators had mutations within or near the MMR-encoding region, which could cause disruptions in the functionality of this system as was suggested for *S. aureus*^51^. It is notable that both mutators belong to two common lineages (CCs 19 and 23) associated with colonization during pregnancy^16^. Within the non-mutator postpartum strains, those belonging to CCs 1 and 23 along with CC-12 had the most SNPs overall. Nevertheless, we cannot rule out the possibility that the prenatal strains have other unique traits that may have made them more prone to mutations in DNA repair systems. Because only two mutators were identified in our study, more *in vitro* experiments and cohort studies are needed to determine how GBS mutators emerge in different populations. GBS strain traits (e.g., serotypes, CCs) are variable across geographic locations^16^ while many countries lack IAP protocols altogether^54^. Therefore, different selection pressures in distinct locations will likely have an impact on the type and number of mutations identified in specific strain populations.

Since a high frequency of nonsynonymous SNPs was detected in the postpartum genomes, which could influence protein function, and multiple strains had mutations in the same genes, it is likely that some genes evolve in parallel under selection, leading to enhanced fitness and similar adaptations^57^. We cannot conclude, however, that the mutations arose solely due to IAP exposure, as we are unable to compare the impact of IAP versus no IAP on genomic changes in this study. IAP was the standard protocol for GBS-positive participants and hence, only three participants with persistent colonization were not given IAP. Although the postpartum strains from these three participants had a slightly lower average mutation rate (1.50 x 10^-6^) than the average of the 19 strains with IAP exposure (9.76 x 10^-6^), the difference was not significant (t-test, p=0.24). Since all participants did not have the same antibiotic exposures, this variation could impact our selection measures and mutation rate estimates. Indeed, not all participants were given the same antibiotic or received the same number of doses, and some reported postpartum antibiotic use prior to the second sampling. We also could not account for other factors (e.g., changes in the vaginal microbiome or host susceptibility, over-the-counter medication use, complications during childbirth, diet, etc.) that may impact selection and evolution *in vivo*. These limitations highlight the need for long-term studies with a larger and more diverse patient population as well as more serially collected samples from participants who receive the same IAP exposure (e.g., concentration, doses and type of antibiotic). This type of sampling scheme could have helped determine whether ancestral intermediaries exist in the evolution of strain 22.2, for instance, which shares an ST with the prenatal 22.1 strain but differs in gene content. Alternatively, it could pinpoint the timing when a new strain type was acquired after childbirth.

In all, this analysis highlights the need for future studies that will examine potential mechanisms that could be linked to persistent colonization and to understand how mutators emerge in populations under selection. Even though the biofilm assays revealed functional differences in strains collected before and after IAP and childbirth, additional experiments are needed to determine if the mutations identified in this study can impact this phenotype or others. The lack of mechanistic confirmation is a limitation of our descriptive study and hence, we cannot confirm a causal relationship between IAP and mutation rates. Larger prospective clinical studies are critically important to determine how IAP and other selective pressures, which are encountered by GBS during childbirth, contribute to mutation rates and persistence *in vivo* and to further define their role in colonization and disease. Despite our findings, it is important to note that the prophylactic use of antibiotics to protect newborns from invasive GBS infections far outweighs the risks imposed by evolutionary events that may enhance colonization.

## METHODS

### Bacterial isolate selection and characterization

A subset of 97 GBS isolates recovered from vaginal-rectal swabs from 58 pregnant participants^23^ were included in this analysis. Isolates were recovered from the same participant during a prenatal (35-37 weeks gestation) visit and during a postpartum visit, which took place six weeks after childbirth^23^. Isolates were previously characterized by MLST, serotyping, and cps typing^22^. Participants with GBS at both samplings were considered to have persistent colonization resulting in 80 strains from 41 participants. Seventeen participants had GBS only at the prenatal sampling and were considered to have lost the pathogen by the postpartum visit.

### Whole-genome sequencing (WGS)

A set of 41 of the 97 isolates was sequenced previously and the raw reads were downloaded from the NCBI Sequence Read Archive (SRA); the remaining 56 isolates were sequenced for this study (**Table S2**). Following DNA extraction of GBS grown overnight in Todd-Hewitt broth (THB) at 37°C + 5% CO_2_ using the E.Z.N.A. (Omega Bio-tek Inc., Norcross, GA, USA) or Wizard HMW (Promega, Madison, WI, USA) kits, libraries were constructed at the Michigan Department of Health and Human Services using the Nextera XT library prep kit (Illumina, San Diego, CA, USA). Sequencing was performed using the MiSeq (Illumina) with 2x250 bp paired end reads. Raw reads were trimmed with Trimmomatic v0.39^55^ using “gentle trimming” parameters to trim adapters and remove sequences with average quality scores <15 or that were <36 nucleotides long. The paired-end reads, or single-end reads for SRR517012 that was sequenced using an unpaired method, were assessed for quality using FastQC v0.11.9 (https://www.bioinformatics.babraham.ac.uk/projects/fastqc/); high-quality genomes were defined as those that failed <10% of quality modules.

Sequences were assembled *de novo* using SPAdes v3.13.1 (kmers 21, 33, 55, 77, 99, 127) with mismatch error correction^56^, and quality checked using QUAST v5.0.2^58^ and MultiQC v1.15^59^ with default parameters. High-quality assemblies, which were considered if the N50 and L50 scores were >15,000 and <50, respectively, were annotated using Prokka v1.14.6 with the – proteins option that allows for the use of a custom database^60^. This database was curated from 25 closed *S. agalactiae* genomes downloaded from NCBI to ensure proper annotations (**Table S9**). STs were assigned using PubMLST^61^, though previously determined STs^22^ were used for three strains with poor assembly quality for one of the seven MLST loci.

### Pangenome and phylogenetic analysis

Roary v3.11.2 was used to create a multiFASTA alignment of core genes from the 92 high-quality assemblies using the -i (blastp of 95%) and -e (PRANK aligner 170427) parameters^62^. Maximum likelihood phylogenies were generated with RAxML v8.2.12 (parameters -m GTRGAMMA, 500 bootstrap replicates)^63^. The core-gene alignment was input into MEGA12,^24^ to construct a phylogeny using the Maximum Likelihood method and Tamura-Nei model^64^ of nucleotide substitutions, which used up to 4 parallel computing threads. The tree with the highest log likelihood was used and the percentage of replicate trees was determined adaptively. The initial tree was selected by choosing the tree with the superior log-likelihood between a Neighbor-Joining (NJ) tree,^27^ which uses a matrix of pairwise distances, and a Maximum Parsimony (MP) tree with the shortest length among 10 MP tree searches.

Phylogenies were visualized and annotated using the Interactive Tree of Life v7 (IToL, https://itol.embl.de/)^28^. Neighbor-net trees were generated in SplitsTree v4.19.0^25^ and the PHI test was used to detect recombination with a window size of 100 and a k of 1; a p-value of <0.05 was considered significant. Analysis of the six MMR loci was performed by manually extracting them from the assemblies using Geneious v2022.2.2 (https://www.geneious.com/) followed by a MUSCLE^31^ nucleotide alignment.

### Gene and nucleotide-level analyses

ARGs and virulence genes were extracted from the 92 assemblies with ABRicate (https://github.com/tseemann/abricate) using five ARG databases (CARD^65^, ARG-annot^66^, MEGARes^67^, Resfinder^68^, NCBI AMRFinderPlus^69^) and the virulence-finder-database (VFDB)^70^, respectively. A gene was considered present if it had ≥10% identity and ≥80% coverage and for the ARGs, if it was detected in two or more databases.

Core gene mutations were extracted from the assemblies of each pair of strains recovered from participants with persistent colonization (n=64, 32 pairs). For each pair, the assembled prenatal genome was used as the reference, while the trimmed reads from the postpartum isolate were mapped against the reference. Each core genome was interrogated for mutations including SNPs, insertions, deletions, MNPs, and complex mutations using Snippy v4.6.0 (https://github.com/tseemann/snippy). For discrepancies in gene presence/absence within a pair of strains, Geneious v2022.2.2 was used to confirm ≥15X read coverage and the up and downstream regions were mapped with respect to a given gene to confirm location and completeness of the gene within the contig. Core gene mutations were validated using a ≥10X read coverage cutoff.

### Biofilm Assays

Biofilm assays were performed in triplicate on 30 pairs of strains from persistently colonized participants (n=60 isolates). Values were normalized to blank media controls as we described^39^ previously with the modification of stagnant growth for 24hrs. The magnitude (fold-change) of the difference in biofilm formation was determined for each postpartum strain relative to each respective prenatal strain.

### Data analysis and statistics

Raw WGS output was managed in Microsoft Excel. R v4.1.2 (https://www.r-project.org/) and RStudio v2022.07.1+554 (https://www.rstudio.com/) was used for importing data (readr v2.1.4), data wrangling (devtools v2.4.5, dplyr v1.1.3, forcats v1.0.0, plyr v1.8.8, tidyr v1.3.0, tidyverse v2.0.0), and visualization (ggplot2 v3.4.3, viridis v0.6.4) (https://cran.r-project.org/). R package factoextra v1.0.7 was used to perform and visualize PCA with 95% confidence ellipses for virulence genes using the prcomp() and fviz_pca() functions (https://cran.r-project.org/). Chi- square tests were performed in Microsoft Excel or Epi Info™ v7 (Centers for Disease Control and Prevention, Atlanta, GA, USA); statistically significant associations were classified at p ≤ 0.05. For unnamed genes that acquired mutations, arbitrary “gene ids” using the GBS_00X notation were assigned for easier reference (**Table S6**). For all SNP-only analyses, SNPs were filtered to exclude those that occurred in rRNA and/or uncharacterized, non-coding regions.

As characterized by Snippy, nonsynonymous SNPs included those classified as “initiator codon variant”, “missense”, “start lost”, “stop lost”, and “stop gained” (nonsense), while synonymous SNPs included those referred to as “stop retained” and “synonymous variant”. Measures of evolutionary selection were calculated based on the nonsynonymous to synonymous SNP ratio (πN/πS) across the genomes, as well as across each gene of interest to discern areas under positive (πN/πS >1), neutral (πN/πS = 1), or negative (πN/πS <1) selection. Calculations for evolutionary selection across genes were limited to genes that acquired four or more SNPs.

Student t-tests were used to analyze differences in biofilm formation between isolate pairs and mutation rates across CCs; p<0.05 was deemed significant.

## Data availability

All datasets are available in the supplementary tables and raw sequencing reads for each genome have been deposited in GenBank under several BioProjects listed in **Table S2** unless they were uploaded previously. R scripts are at: https://github.com/macyEpell and via Zenodo (https://doi.org/10.5281/zenodo.16413547).

## Ethics statement

The parent study was conducted in November 1998-April 2000 and involved obtaining informed consent from each patient prior to enrollment. It was approved by the Conjoint Medical Ethics Committee of the University of Calgary and the Ethics Board of the participating University of Alberta Hospitals.

## Acknowledgements

We thank the Research Technology Support Facility and Michigan Department of Agriculture and Rural Development for DNA sequencing, as well as the National Center for *Streptococcus* (Edmonton, Alberta), particularly Marguerite Lovgren, for strain collection and serotyping.

Financial support was provided by the Michigan State University (MSU) Research Foundation (to SDM) and in part, by the Michigan Sequencing and Academic Partnerships for Public Health Innovation and Response (MI-SAPPHIRE) initiative at the MDHHS via CDC through the Epidemiology and Laboratory Capacity for Prevention and Control of Emerging Infectious Diseases Enhancing Detection Expansion program (6NU50CK000510-02-07; to HMB and SDM). Salary support was provided by the National Institutes of Health (R01HD090061 to JAG and SDM, R01AI134036 to JAG), Department of Veterans Affairs Merit Award I01BX005352 (Office of Research) to JAG, and the U.S. Department of Agriculture (2019-67017-29112) to SDM. Student support was provided to MEP by the MSU Department of MGI via the Philipp and Vera Gerhardt Travel and Ralph Evans Awards and the MSU College of Natural Science via two Outstanding Scholar Fellowships.

## Author contributions

Conceptualization: MEP, SDM; methodology: HMB, MEP; validation and formal analysis: MEP, SDM; resources: SDM, HMB; supervision and insight: SDM, HMB, JAG, HDD; data curation: MEP, SDM; writing—original draft preparation, MEP; writing—review and editing and data visualization: MEP, SDM; funding acquisition: SDM, HMB. All authors have read and agreed to the final version of the manuscript.

