## Supplemental figures for "Genomic adaptation in group B *Streptococcus* following intrapartum antibiotic prophylaxis and childbirth"

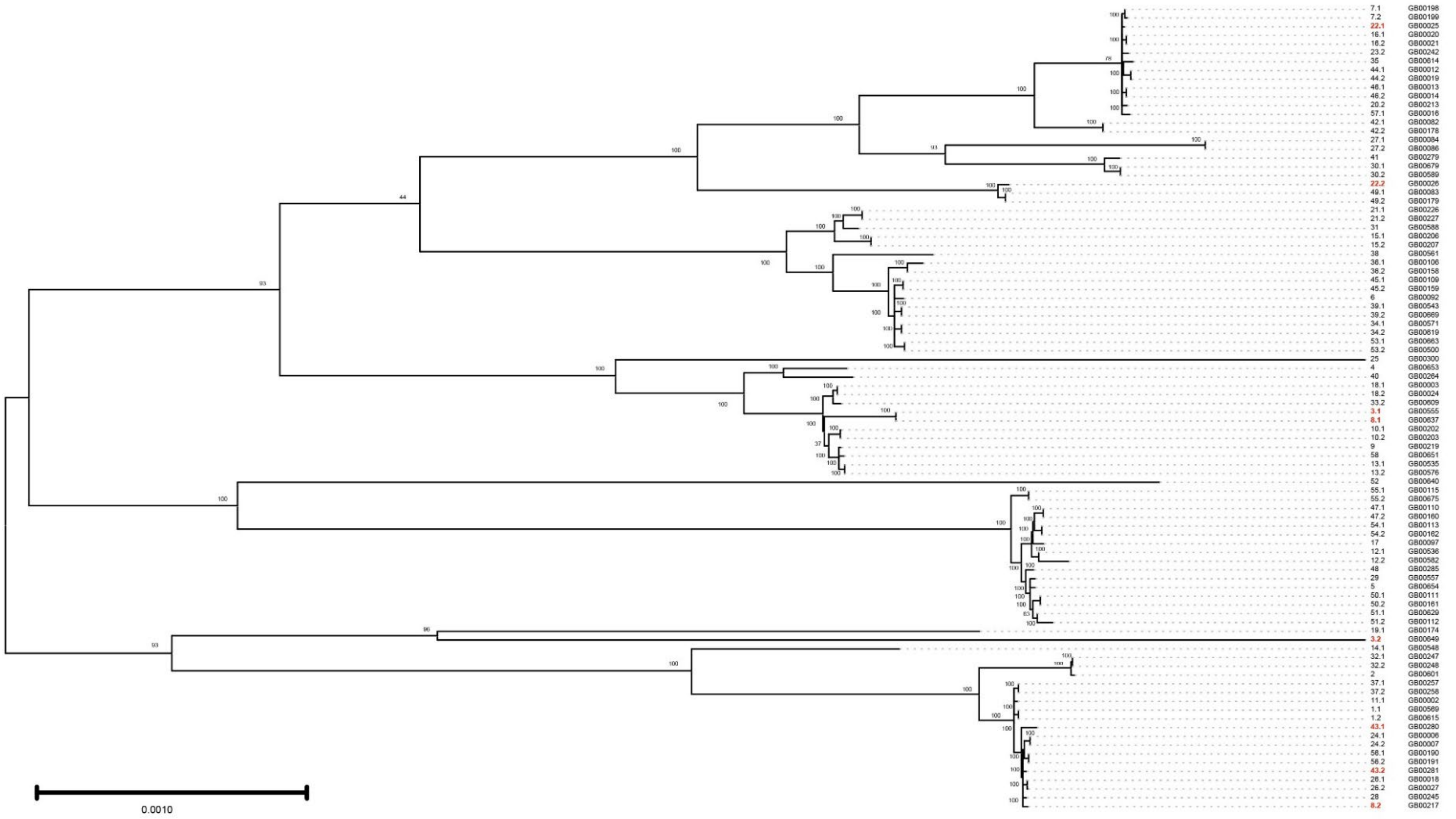


**Figure S1. Maximum likelihood phylogeny showing evolutionary relationships between 92 group B streptococcal strains from individuals before and after childbirth and intrapartum antibiotic prophylaxis.** The phylogeny shows the tree with the highest log likelihood based on 1,368 core genes (1,268,723 positions) inferred using the Maximum Likelihood method and Tamura-Nei model of nucleotide substitutions. The numbers on the branches represent the bootstrap support values, or the percentage of replicate trees in which the associated taxa clustered together out of 88 replicates. The initial tree was selected by choosing the tree with the superior log-likelihood between a Neighbor-Joining tree, which used pairwise distances, and a Maximum Parsimony tree. The tree was generated in MEGA12 using up to 4 parallel computing threads. The participant ID and strain names are shown at the right, with bold red font highlighting the four strain pairs (n=8 strains) that did not group together by pair.

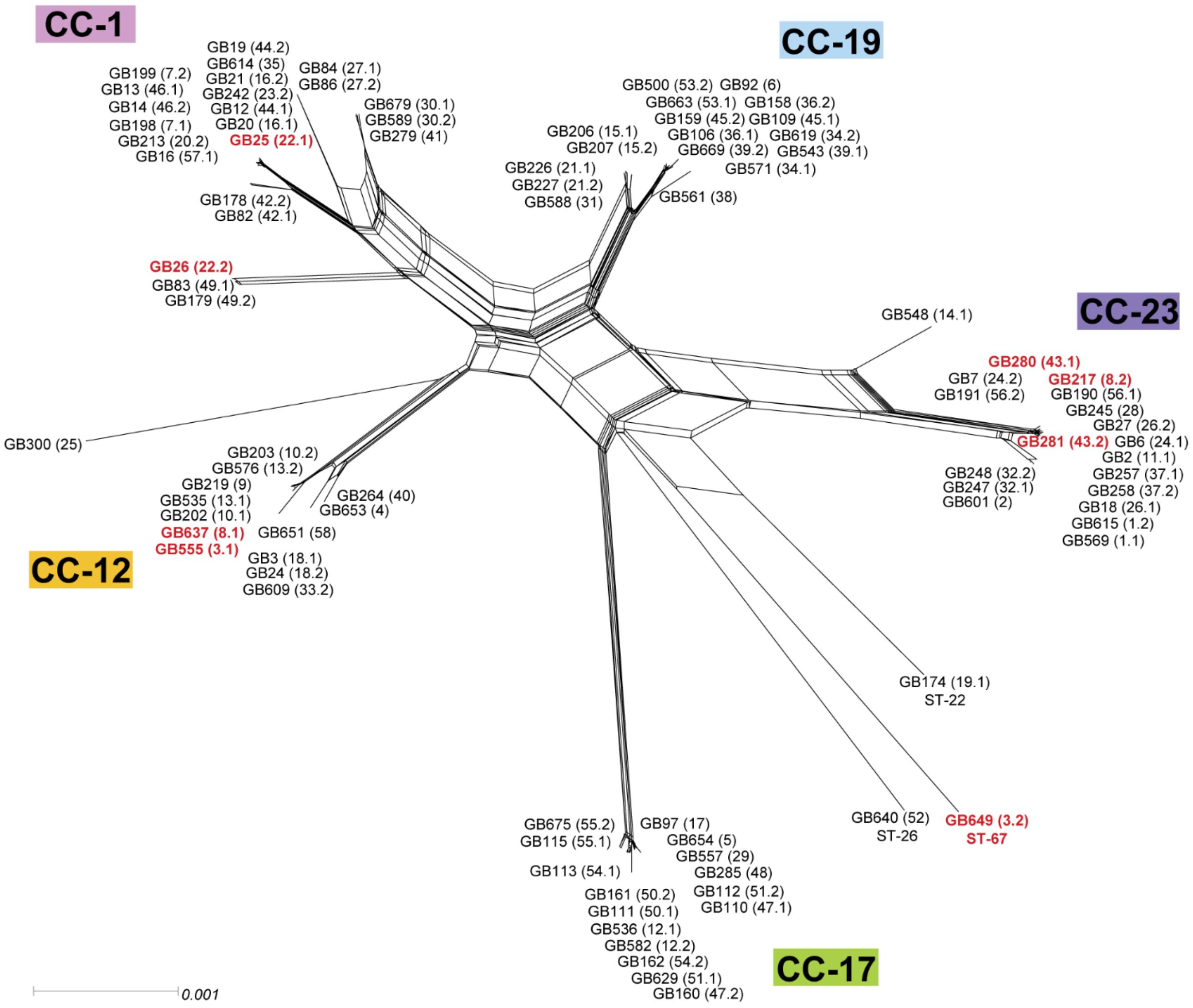


**Figure S2. Neighbor-net tree based on 17,825 parsimony-informative (PI) sites in 1,368 core genes frp, 92 group B streptococcal genomes.** The tree was constructed using SplitsTree v.4.19.0 with a window-size of 100 with k as 1. The pairwise homoplasy index (PHI) revealed significant evidence for recombination (p<0.001). All GBS genomes were included in this analysis, including those from persistently colonized participants who were missing a paired prenatal or postpartum genome due to loss or poor sequence quality. The paired strains for four participants (3,8, 22, and 43) that did not group together are shown in bold red font.

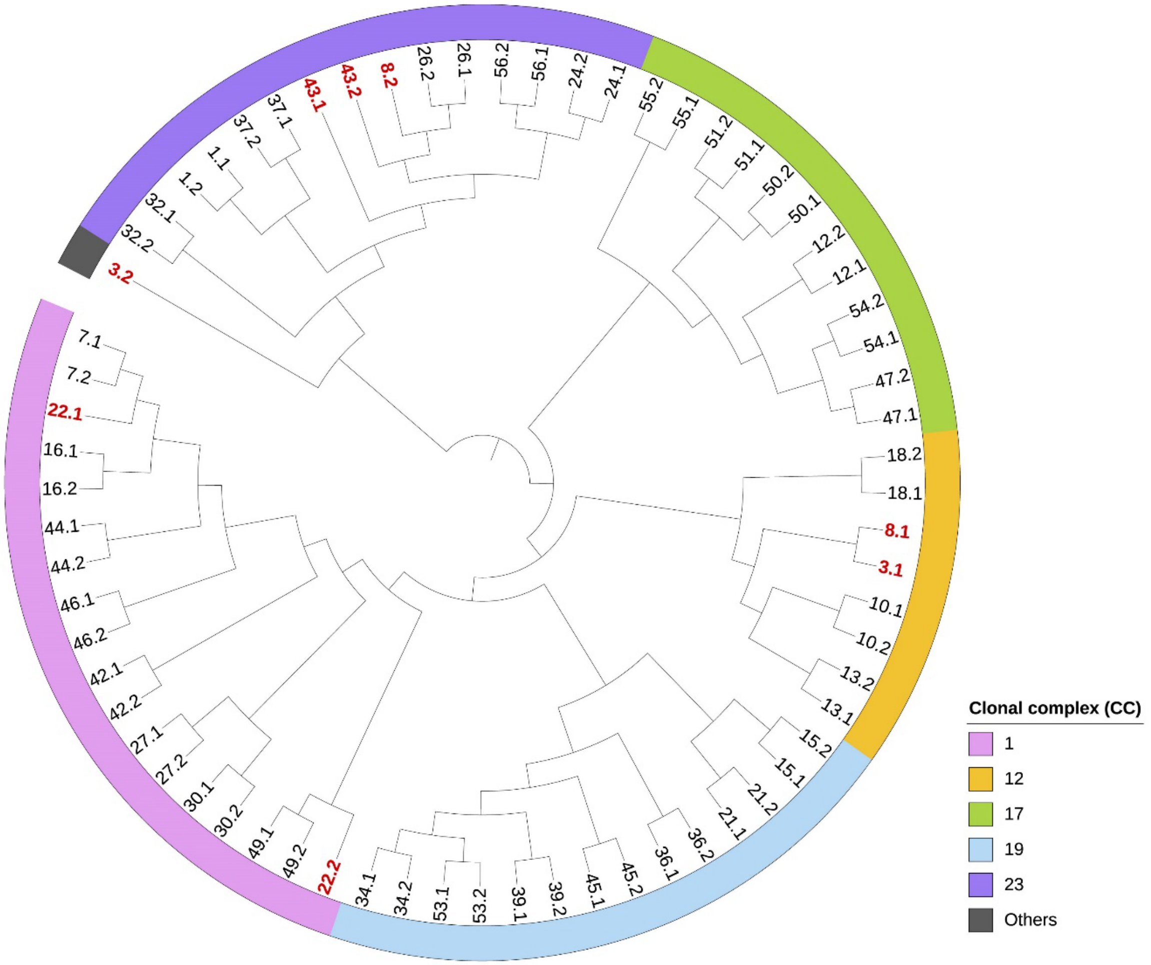


**Figure S3. Core gene phylogeny showing only the 68 paired strains from participants before and after IAP and childbirth by clonal complex (CC).** The bootstrap consensus tree is rooted at the midpoint and was constructed using 1,368 core genes (1,268,723 positions). It was inferred from 88 replicates with the collapsed branches corresponding to partitions reproduced in less than 50% of replicate trees. The original tree (Figure S1) was chosen to represent the one with the highest log-likelihood between a Neighbor-Joining (NJ) tree and Maximum Parsimony (MP) tree and was generated in MEGA12 and visualized with iToL v6 after rooting at the midpoint. Participant strain IDs are at the end of each branch with a ".1" representing the prenatal strains and a ".2" representing the postpartum strains. IDs in bold red font indicate the paired prenatal and postpartum strains from four participants (IDs 3, 8, 22, and 43) that did not cluster together in Figures 1, S1, and S2.

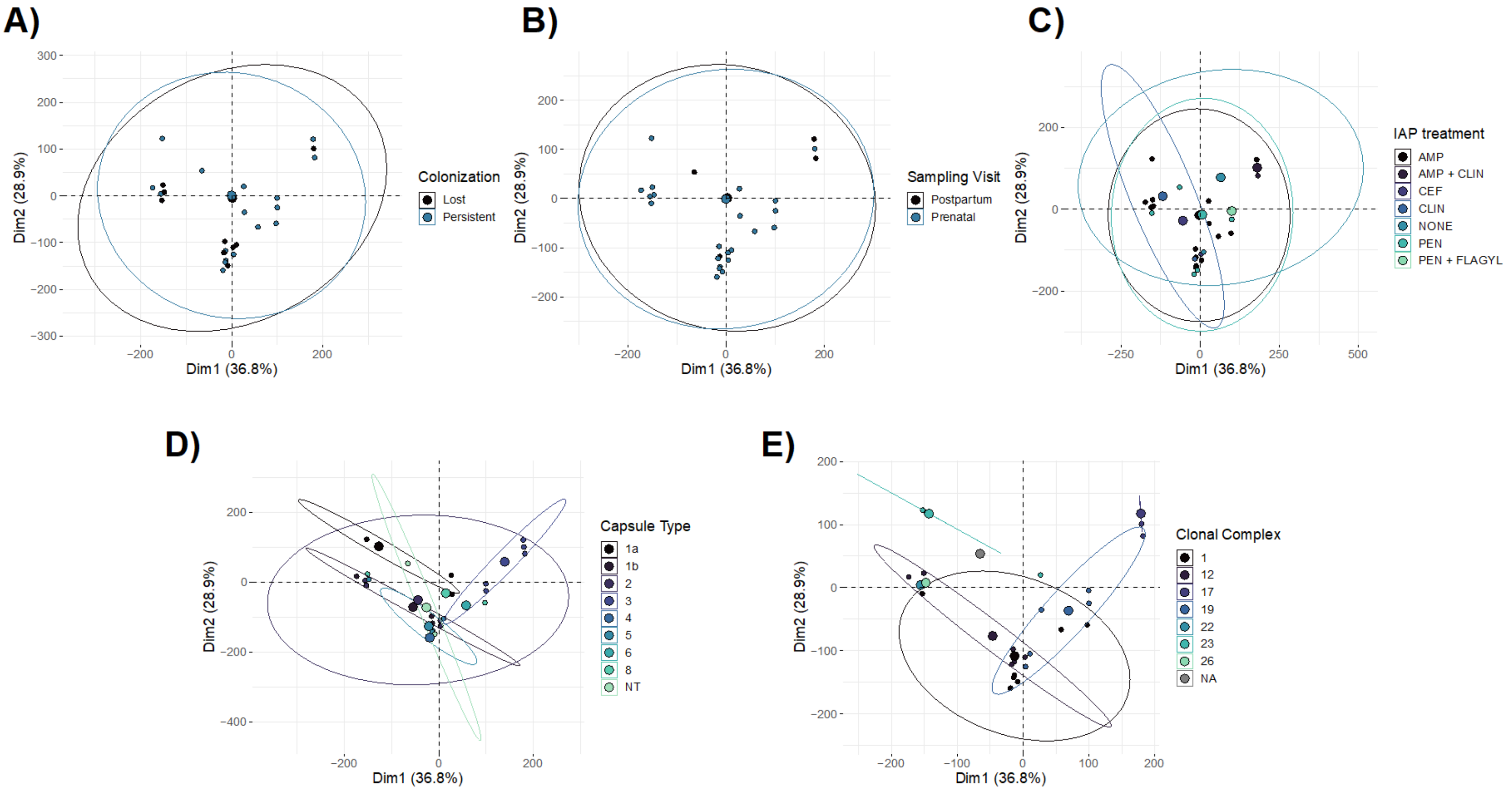


**Figure S4. Principal component analysis (PCA) of virulence gene profiles.** Percent identity values of all virulence genes detected in the 92 GBS genomes were used for the PCA. In each plot, the single PCA was stratified by **A**) colonization status (lost versus persistent), **B)** sampling visit, **C**) IAP treatment, **D**) capsule type, and **E**) clonal complex. The first principal component is shown on the x-axis (Dim1) and the second (Dim2) is on the y-axis, representing 36.8% and 28.9% of the variation in the sample set, respectively. Each point represents a strain's virulence gene profile (i.e., the set of virulence genes that are present or absent). Ellipses indicate the 95% confidence interval of relatedness and the larger points in each plot represent the cluster's median in which the ellipse is drawn. For panels that have categories with too few points to calculate a confidence ellipse (C-E), these categorical clusters are represented with their median value without an ellipse. AMP=ampicillin; CLIN=clindamycin; CEF=cefazolin; PEN=penicillin.

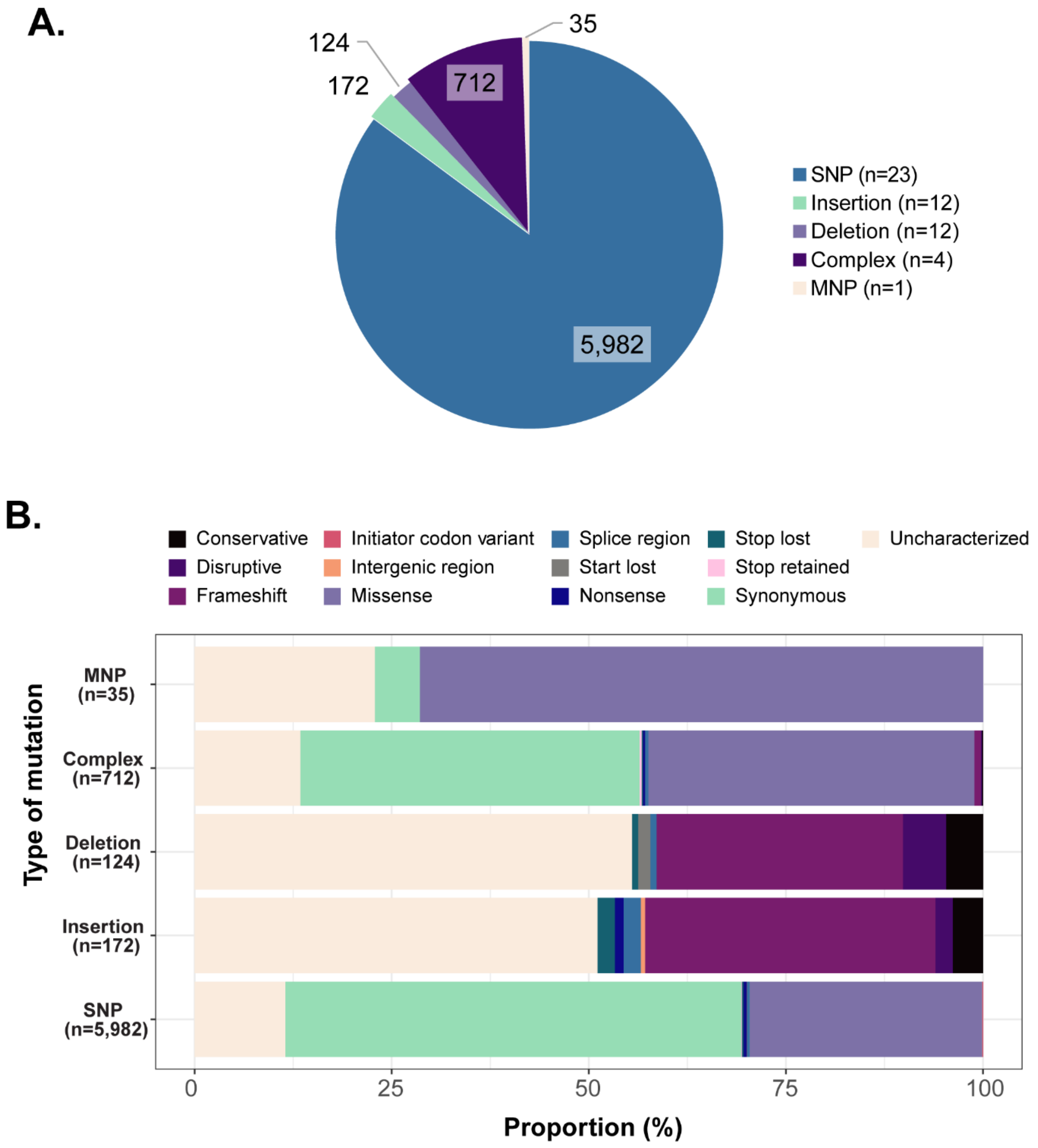


**Figure S5: Type of core gene mutations detected in the postpartum GBS genomes from participants following IAP. A**) Distribution of the type of mutations (n=7,025) found in the 24 postpartum genomes. The number of postpartum genomes with each mutation is shown in parentheses in the legend. **B)** Boxplot displays the proportion (x-axis) of each mutation outcome (colors) for each mutation category (y-axis) with raw mutation counts noted in parentheses. Nonsynonymous SNPs included those classified as “initiator codon variant”, “missense”, “start lost”, “stop lost”, and “stop gained” (nonsense), while synonymous SNPs included those referred to as “stop retained” and “synonymous variant”. MNP= multiple nucleotide polymorphisms

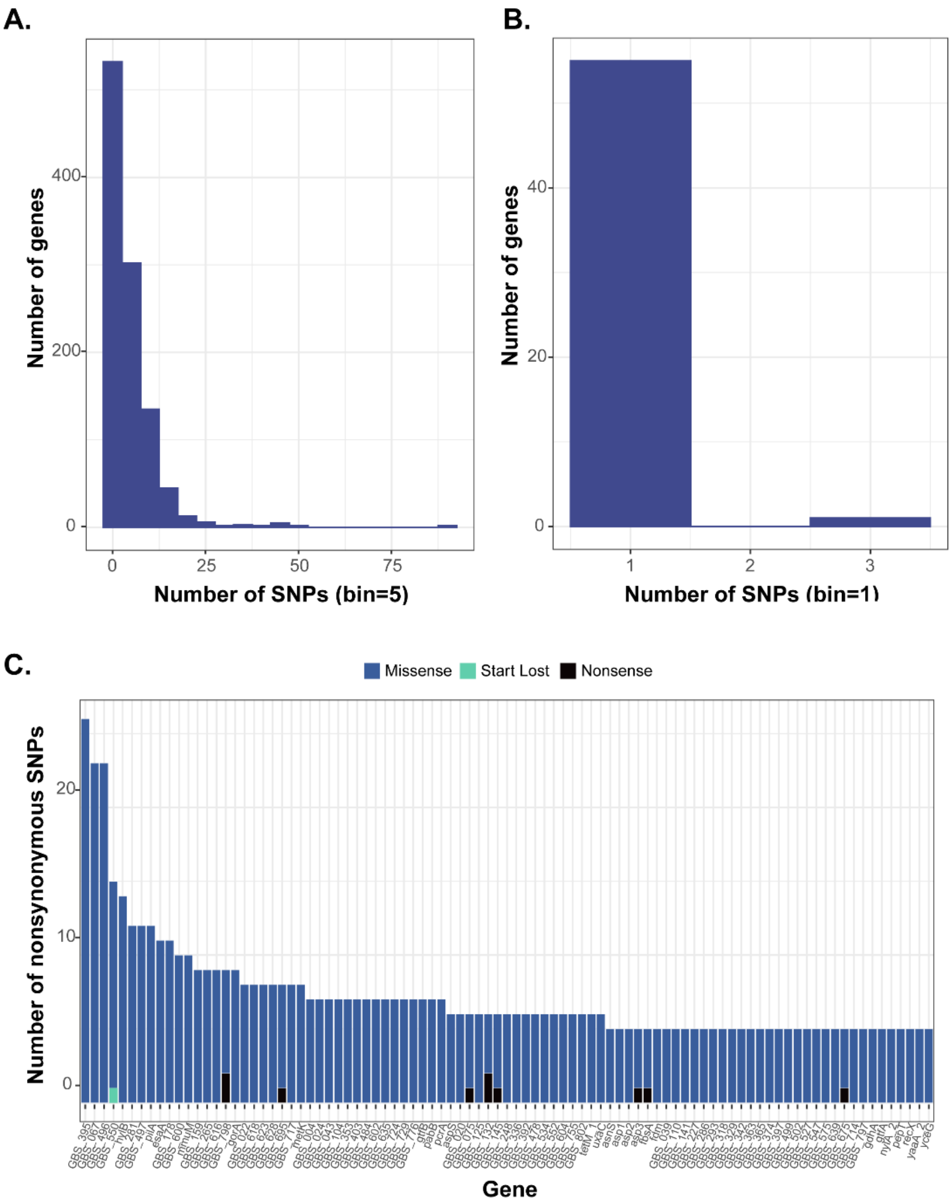


**Figure S6. Number and distribution of SNPs detected within genes from the 32 postpartum genomes relative to the respective prenatal genomes.** The distribution of SNPs (x-axis) by the number of genes (y-axis) in the **A)** three outlier genomes with the most mutations, using a bin width, or interval size, of 5, and **B)** the remaining genomes, using a bin width, or interval size, of 1. **C)** The distribution of nonsynonymous SNP types (missense=blue, start/lost=green, nonsense=black) is shown for the three outlier genomes by gene (x-axis) for the 91 genes with >4 nonsynonymous SNPs.

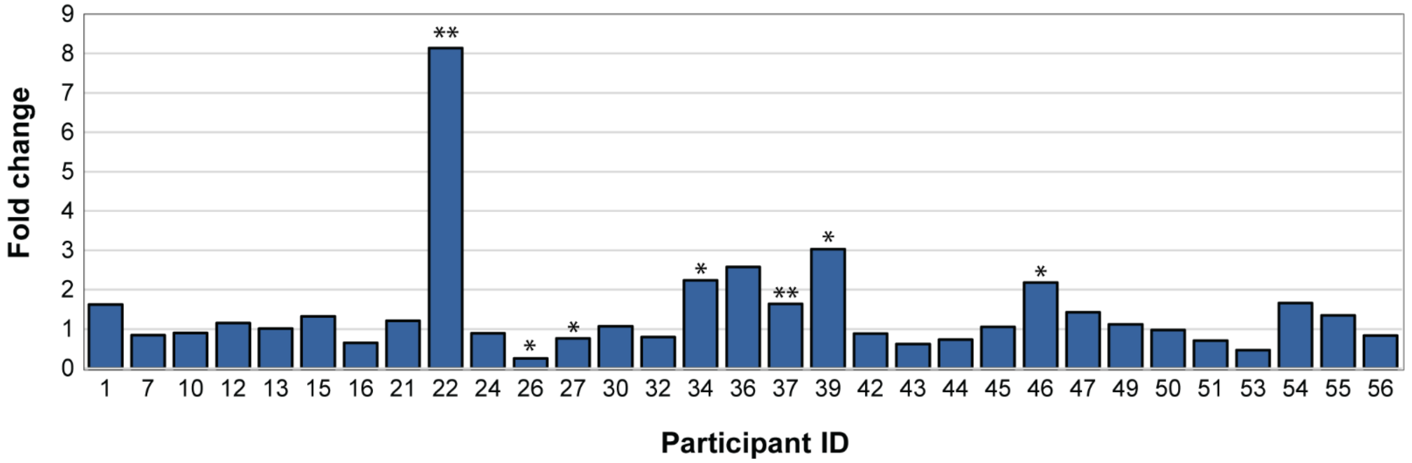


**Figure S7. Fold change difference in biofilm formation between 62 paired persistent isolates with the same ST.** Based on the absorbance values ($OD_{595}$), the fold change (y-axis) in biofilm formation was calculated between the pair of isolates from each participant (x-axis). The average absorbance value (n=3 replicates) for the postpartum isolate was divided by the average absorbance of its respective prenatal isolate (n=3 replicates). A fold change >1 indicates an increase in biofilm formation in the postpartum isolate relative to its prenatal isolate, while a fold change <1 indicates a decrease in biofilm formation. One-tailed, paired t-tests were calculated to assess significant differences in biofilm formation between isolate pairs ($*=p<0.05$, $**=p<0.01$).
